# Radiotherapy-induced vascular dysfunction alters neutrophil phenotype to promote tumor cell colonization of mammary adipose tissue

**DOI:** 10.64898/2026.10.03.756445

**Authors:** Shannon E. Martello, Bridget H. Stevenson, Holden Korbey, Kevin C. Corn, Tian Zhu, Erica J. Lin, Marissa Paul, Kirsten E. Stubenrauch, Jonathan Wang, Alessandra Perez, Marjan Rafat

**Affiliations:** Department of Chemical and Biomolecular Engineering, Vanderbilt University, Nashville, TN, USA; Department of Biomedical Engineering, Vanderbilt University, Nashville, TN, USA; Department of Biological Sciences, Vanderbilt University, Nashville, TN, USA; Department of Chemistry, Susquehanna University, Susquehanna, PA, USA; School for Science and Math at Vanderbilt, Vanderbilt University, Nashville, TN, USA; Department of Radiation Oncology, Vanderbilt University Medical Center, Nashville, TN, USA; Vanderbilt-Ingram Cancer Center, Vanderbilt University Medical Center, Nashville, TN, USA

**Author notes:** Author for correspondence: Marjan Rafat, PhD, Engineering and Science Building, Rm. 426, Vanderbilt University, Nashville, TN 37212.

## Abstract

Recurrence remains a significant clinical challenge for triple-negative breast cancer patients and radiotherapy (RT) is a key therapeutic tool that significantly reduces the risk of relapse. Despite this, patients with chronic lymphopenia and a high neutrophil:lymphocyte ratio (NLR) experience recurrence at significantly higher rates than those whose hematological biomarkers return to baseline post-RT, and there are currently no treatments to prevent this relapse. Here, we utilized *in vivo* mouse models of lymphocyte-deficiency to identify the mechanism behind systemic immune status and recurrence. Using a model of RT-induced circulating tumor cell (CTC) colonization, we found that neutrophils are sustained in the irradiated, lymphocyte-deficient mammary tissue up to 10 days post-RT. Persistent neutrophil accumulation was accompanied by excessive vascular remodeling and tumor cell colonization of the mammary tissue, both of which were abated upon systemic neutrophil depletion. Neutrophils in the colonization-permissive microenvironment exhibited a state of pathological activation consistent with a polymorphonuclear myeloid-derived suppressor cell (PMN-MDSC) phenotype that was not seen in systemic compartments, highlighting a microenvironment-induced phenotype shift. This phenotype was recapitulated *in vitro* when naïve neutrophils were exposed to the secretome of RT-induced senescent endothelial cells (ECs). We found that these activated neutrophils then regulated the dysfunction of irradiated ECs in a canonical NF-kB-dependent manner, increasing their barrier permeability and activating downstream inflammatory signals. Our findings not only establish neutrophils as a critical component of tumor cell colonization in lymphocyte-deficient subjects but also uncover a positive feedback loop between neutrophils and irradiated vasculature that leads to aberrant tissue remodeling and ultimately tumor cell colonization. This work deepens our understanding of how systemic factors influence treatment-damaged microenvironments, which could ultimately lead to identification of molecular targets for prevention of RT failure.

## Introduction

Most breast cancer patients will receive radiotherapy (RT) due to its demonstrated ability to reduce relapse^1^, yet nearly one in three patients with triple-negative breast cancer (TNBC) will experience recurrence within five years of diagnosis^2^. Surgery and hematological parameters strongly dictate outcomes for patients^3–5^. For TNBC patients that receive RT, a sustained low absolute lymphocyte count (ALC) and a high neutrophil:lymphocyte ratio (NLR) in the months following RT is associated with a high risk for local recurrence^6,7^, which is often accompanied by distant metastasis. We have previously shown in mouse models of lymphocyte-deficiency that RT-damaged mammary adipose tissue is colonized by tumor cells and characterized by persistent macrophage infiltration in the absence of T lymphocytes^6,8,9^, demonstrating potential immune-mediated recurrence mechanisms. Still, the cause of this sustained immune cell infiltration and the mechanistic link between high NLR and recurrence remain unclear.

Neutrophils have recently been established as key regulators of metastasis and chronic inflammation, functioning well beyond their conventional acute response^10–12^. Neutrophils activated by damaged tissue environments—whether from environmental stressors or direct therapeutic treatment—are capable of both direct and indirect regulation of tumor cell colonization. For example, the release of neutrophils extracellular traps (NETs) can sequester cancer cells from circulation^13^, alter the molecular signaling of circulating tumor cells (CTCs)^14^, and create blockages in the vasculature that induce necrosis^15^. It has also been shown that irradiated lung tissue prolongs neutrophil retention and promotes their degranulation to facilitate metastasis^16^. While these studies strongly support the role of neutrophils in tissue remodeling and pre-metastatic niche formation, most studies focus on the lung or tumor microenvironment, leaving the effects of chronic neutrophil infiltration on the normal breast microenvironment understudied. Furthermore, the impact of systemic factors like low ALC on neutrophil interactions with locally damaged microenvironments is currently unknown.

In the current study, we utilized lymphocyte-deficient mouse models of RT-induced CTC colonization to identify the effects of systemic lymphopenia on the irradiated mammary microenvironment and determine the role of neutrophils in local recurrence. We characterized neutrophil infiltration post-RT and identified associated alterations to the neutrophil phenotype and vasculature that coincide with tumor cell colonization. Mechanistically, we hypothesized that RT induces a persistent senescent, inflammatory phenotype in endothelial cells that initiates a reciprocal feedback loop with neutrophils to prolong tissue damage and facilitate formation of a pro-recurrent niche. Importantly, these findings were only observed in mice with T lymphocyte reduction, warranting further study of recurrence mechanisms in models with clinically relevant systemic states to aid development of interventions for high-risk populations.

## Materials and Methods

### Cell culture

Luciferase-labeled 4T1 mouse mammary carcinoma cells (4T1^luc^) were obtained from ATCC and maintained in RPMI-1640 supplemented with 10% fetal bovine serum (FBS) and 1% penicillin/streptomycin (P/S). Murine embryonic endothelial cells (ECs) (C166; ATCC, CRL-2581) were cultured in DMEM supplemented with 10% FBS and 1% P/S. For neutrophil co-culture experiments, murine neutrophils were differentiated from the murine promyelocyte cell line (MPRO 2.1; ATCC, CRL-3604) by daily supplementation of base media (IMDM with 1% P/S, 20% horse serum, and 10ng/mL granulocyte-macrophage colony-stimulating factor (GM-CSF)) with 10µM all-trans retinoic acid (ATRA; Sigma) for 72 hours. Neutrophils were then used in experiments by day 4 of differentiation. All cells were cultured under 5% CO_2_ at 37°C in a humidified incubator. All cell lines were tested monthly for mycoplasma contamination using a MycoAlert Mycoplasma Detection Kit (Lonza).

### In vivo mouse models

Animal studies were performed in accordance with the institutional guidelines and protocols approved by the Vanderbilt University Institutional Animal Care and Use Committee. All studies were performed on age, sex, and strain-matched animals treated at the same time of day, according to ARRIVE 2.0 guidelines^17^. Female BALB/c or athymic Nu/Nu mice (8-10 weeks old) were inoculated with 5x10^4^ 4T1^luc^ cells into the right #4 mammary fat pad (MFP). For CD8^+^ T cell-reduction experiments, mice were injected intraperitoneally (i.p.) with 500µg of anti-CD8a [2.43] (BioXcell) or anti-IgG2b isotype control [LTF-2] (BioXcell) at the time of tumor cell inoculation and every 5 days thereafter. For Ly6G depletion experiments, mice were injected i.p. every other day with 100µg of anti-Ly6G [1A8] (BioXcell) or anti-IgG2a isotype control [2A3]. Confirmation of both CD8^+^ T cell depletion and Ly6G^+^ cell depletion was confirmed by analyzing spleens for the respective cell type via flow cytometry. Tumor length and width were measured using digital calipers starting one week post-inoculation. Mice were euthanized in a carbon dioxide chamber for post-mortem excision of tissues for histologic and flow cytometry analysis or for bone marrow-derived neutrophil isolation, which is a method approved by the Panel on Euthanasia of the American Veterinary Medical Association and is considered humane. Cervical dislocation was performed subsequently as a secondary physical method as required by the NIH Office of Laboratory Animal Welfare

### In vivo irradiation

When tumors reached 100mm^3^, the contralateral #4 MFP was irradiated to 20 Gy using a 300 kVp cabinet x-ray system filtered with 1.57 mm aluminum. During irradiation, mice were anesthetized with 2.5% isoflurane and shielded using a 2 cm thick Cerrobend shield with 1 cm wide and 1.5 cm long apertures to expose the tumors. Transmission through the shield was less than 1%.

### Immunohistochemistry

At 10 days post-RT, MFPs were harvested and placed in 10% neutral buffered formalin (NBF) for 24h at 4°C. Tissues were then rinsed and kept in 70% ethanol until processing. Tissues were embedded in paraffin and sectioned (4µm) onto slides. Sections were deparaffinized, rehydrated, and incubated in citric acid buffer (pH 6) for 30 min at 95°C for antigen retrieval. Slides for MPO/CitH3 staining were incubated in Tris-EDTA buffer (pH 9). For chromogenic immunohistochemistry (IHC), sections were then treated with 3% hydrogen peroxide in methanol and incubated with avidin and biotin before blocking with PBS/10% goat serum/1% BSA/22.5 mg/mL glycine for 1 hour at room temperature. Primary antibody incubation was performed overnight using Ly6G primary antibodies followed by sequential incubation with horseradish peroxidase-conjugated streptavidin, biotinylated secondary antibodies, and substrate from a DAB peroxidase substrate kit (Vector Labs). Sections were counterstained with hematoxylin 7211. For immunofluorescence (IF) detection, blocking was performed after antigen retrieval using the above blocking solution with donkey serum (CD31/VEGF) or goat serum (all other antibodies). Primary antibody incubation was performed overnight at 4°C with the following antibodies: CD31, VEGF, F4/80, myeloperoxidase (MPO), citrullinated histone H3 (CitH3), S100A9, and p21. Secondary antibody incubation was then performed for 1 hour at room temperature, followed by autofluorescence quenching with TrueBlack. Nuclear staining with 4’,6-diamidino-2-phenylindole dihydrochloride (DAPI) was then performed and sections were mounted with a coverslip. A no primary control was included for all conditions in both detection methods of IHC.

### IHC analysis

All slides were imaged using an inverted Leica microscope (DMi8) using a Leica DFC9000GT sCMOS fluorescence or MC190 HD digital camera. For each section, at least 30 representative images were taken and Fiji^18^ was used to process and analyze all images. For each staining analysis, 10 images were then randomly selected using a random number generator and downstream analysis was performed. For Ly6G, F4/80, and S100A9 staining, cells per field were hand counted using Fiji’s Cell Counter plugin. For vessel analysis, the area of positive CD31 staining was first identified based on the no primary control sample. Vessel counts per mm^2^ were then determined in Fiji by setting a threshold to measure only areas of positive staining, filling the vessel lumens to avoid double-counting, and quantifying the number of vessels per field using the Fiji Analyze Particles function. CD31+ area was then divided by the image area to generate the number of vessels per area. For CD31-VEGF co-staining analysis, a mask of CD31+ area was created and overlaid onto each corresponding VEGF image. The integrated density of VEGF within each vessel was then measured and averaged per field.

### Flow cytometry

For *in vivo* experiments, blood was drawn retro-orbitally prior to sacrificing. MFP, bone marrow, and spleens were then harvested for downstream processing into a single cell suspension before staining. Briefly, blood was subject to ACK lysis and counted. MFPs were digested in collagenase II for 45 minutes at 37°C under constant agitation and filtered through a 100µm strainer. Bone marrow was isolated according to previously established protocols^19^ and red blood cells were lysed with ACK lysis buffer. Spleens were filtered through a 40µm strainer and red blood cells were lysed with ACK lysis buffer. All cell suspensions were then counted. At least 1x10^6^ cells were stained with Aqua fixable viability (Invitrogen) and FC receptors were blocked using anti-CD16/32 (BioLegend) while simultaneously staining for all surface markers. Cells were incubated in the dark at 4°C, rinsed, and fixed in 1% NBF for 20 minutes. For intracellular markers, cells were permeabilized for 5 minutes using a 1X permeabilization buffer (eBioscience) and stained with antibodies in the 1X buffer for 30 minutes at room temperature. Subsequent washes were performed in the permeabilization buffer and samples were analyzed on a three-laser Amnis CellStream machine (Cytek). FlowJo software (Becton Dickinson) was used for analysis. The following antibodies were used for analysis: CD45-BV650 [30-F11] (1:200; BioLegend 103151), CD11b-APC [M1/70] (1:200; eBioscience 17-0112-82), Ly6G-efluor450 [1A8] (1:100; eBioscience 48966882), Ly6C-APC-efluor780 [HK1.4] (1:200; eBioscience 47-5932-82), CD84-PE [mCD84.7] (1:200; BioLegend, 50-164-046), COX2-AF647 [D5H5] (1:50; Cell Signaling 81707), and ARG1-AF488 [A1exF5] (1:50; eBioscience 53-3697-82).

### Bulk RNA-sequencing

Total RNA was extracted from MFPs and processed for downstream sequencing analysis by the Vanderbilt Technologies for Advanced Genomics (VANTAGE) core facility. For RNA extraction, a hybrid method utilizing Trizol reagent and Qiagen RNeasy kit was used. The tissue samples were homogenized in the presence of 1ml Trizol and 1.5mm stainless steel bead, using the Qiagen TissueLyser II instrument for 2min at 25Hz, twice. Homogenized samples were incubated for 5 min at room temperature after which 200ml. Chloroform was added, mixed well and incubated for 3min at room temperature. Samples were centrifuged at 10,000 g at 4C after which the aqueous layer was transferred to a new tube and mixed with equal volume of 70%. The samples were loaded to Qiagen RNAeasy column, and the samples were processed according to the protocol, including the optional on column DNase treatment. RNA was eluted in 35ul of RNase free water.

For sequencing, the Illumina NovaSeq 6000 platform with multiplexed paired-end 150 bp reads was used by VANTAGE. Processing of bulk RNA-seq data was then performed by Vanderbilt’s Creative Data Solutions by aligning to the reference genome and performing subsequent quality control analyses to determine sequencing accuracy and data integrity. Principal component analysis was applied for quality assessment and identification of outliers and/or batch effects. Reactome software was then used for downstream biological analyses and determination of differentially expressed genes and predicted pathway activation.

### RT-qPCR

MFPs were resected and immediately placed in 1 mL of RNAlater solution (Qiagen). Tissues were then incubated for at least 24h at 4°C under constant agitation before RNAlater was removed and tissues were stored at -80°C until further use. To homogenize samples, MFPs were bathed in liquid nitrogen and then ground into a fine powder using a mortar and pestle. The powder was then lysed in Trizol using a handheld homogenizer and lysates were stored at - 80°C at least overnight. For ECs and neutrophils, cells were washed with ice cold PBS, lysed in Trizol, and stored at -80°C at least overnight. Lysates were thawed at room temperature under constant agitation before passing through a QIAshredder spin column and proceeding with RNA isolation using the PureLink Mini Kit (Invitrogen) according to the manufacturer’s protocol. For qPCR, at least 500ng of RNA was used to synthesize cDNA using the High Capacity cDNA Reverse Transcription Kit (applied Biosystems) according to the manufacturer’s protocol. cDNA was diluted 1:5 using ultra-pure water. Individual qPCR reactions were performed using PowerUp SYBR Green Master Mix (Applied Biosystems) and Custom DNA Oligos (Sigma). Reactions were run using a QuantStudio 5 (Applied Biosystems) and genes were normalized to *Actb*.

### In vitro endothelial cell irradiation

C166 ECs were grown to 70% confluency in T75 flasks and irradiated to a dose of 10 Gy using a cesium source. Following irradiation, cells were immediately passaged into new plates or flasks with reduced serum (2%) to prevent overgrowth at longer timepoints and enable optimal detection of secreted cytokines in the conditioned media. Cell counts at each time point post-RT were kept consistent between control and irradiated cells.

### 5-Ethynyl-2’-Deoxyuridine (EdU) Incorporation Assay

EdU incorporation was measured using the Click-iT Plus EdU Imaging Kit (Invitrogen) following the manufacturer’s protocol. Briefly, ECs were irradiated, passaged onto chamber slides, and cultured in 2% FBS media until the indicated timepoint post-RT. At each timepoint, cells were incubated with 10µM EdU for 1h before fixing in 10% NBF for 10 minutes. Cells were then washed and permeabilized in 0.5% Triton X-100 before incubating in the supplied reaction cocktail for 30 minutes at room temperature. Cells were counterstained with DAPI and mounted with a coverslip. Images were acquired with a DMi8 inverted microscope (Leica) and the fraction of EdU^+^ cells was quantified using Fiji.

### Immunocytochemistry and p21 analysis

Cells were grown on chamber slides and at the indicated timepoint post-RT were rinsed in PBS and fixed for 10 minutes at room temperature in 10% NBF. Cells were then permeabilized in 0.1% Triton X-100, rinsed, and blocked in PBS/10% goat serum for 1 hour at room temperature followed by overnight incubation with anti-p21 (Cell Signaling) at 4°C. Secondary antibodies were incubated with Phalloidin for 1 hour at room temperature before counterstaining with DAPI and mounting with a cover slip. At least 10 representative images were taken per sample using a DMi8 inverted microscope (Leica). Expression of p21 was quantified using Fiji by setting a minimum threshold for positive staining with a no primary control sample. Percent positive area was then measured using the analyze particles function.

### Senescence-associated beta-galactosidase (SA-β-gal) assay

SA-β-gal activity was measured using the Senescence beta-Galactosidase Staining Kit (Cell Signaling) following the manufacturer’s protocol. Briefly, ECs were fixed at room temperature for 10 min in 1X fixative (Cell Signaling) followed by three DPBS rinses before incubating in X-gal staining solution (pH 6.0) at 37°C for 24 hr. Representative brightfield images were acquired via a DMi8 inverted microscope (Leica).

### Conditioned media collection

At the indicated timepoint post-RT, EC media was collected and filtered through a 0.45µm syringe filter to remove cell debris. Filtered samples were then aliquoted and stored at -80°C until use for profiling or neutrophil experiments.

### Cytokine profiling

Conditioned media from three independent irradiations was collected at 2 and 7 days post-RT from 0 and 10 Gy samples. Samples were then analyzed for changes in cytokine secretion via the mouse 32-plex Affymetrix kit, which was processes by Eve Technologies Corporation (Calgary, Alberta, Canada).

### Bone marrow-derived neutrophil isolation

Bone marrow was extracted from the femurs and tibias of naïve female BALB/c mice (6-8 weeks old) as detailed above. Magnetic-activated cell sorting (MACS) was then performed using a mouse neutrophil isolation kit (Miltenyi) to negatively select for Ly6G^+^ cells. Purity of the isolated population for Ly6G^+^Ly6C^mid^ cells was confirmed via flow cytometry. Cells were used immediately in downstream assays.

### Analysis of BMDNs in conditioned media

Isolated BMDNs were resuspended in EC conditioned media and plated at a concentration of 2x10^6^ cells/mL in a U-bottom 96-well plate. Cells were incubated for 24 hours for phenotypic analysis (flow cytometry, gene expression). Cells were then pelleted and stained for flow cytometry or lysed in Trizol for RNA isolation as detailed above.

### NETosis assay

SYTOX green (Invitrogen) has been demonstrated to correlate well with release of neutrophil extracellular traps^20^ and was therefore used in BMDN conditioned media experiments to measure capacity for NETosis. BMDNs were resuspended in EC conditioned media as above and SYTOX green was added at a final concentration of 1:50,000. Cells were then incubated for 2 hours to allow for activation and NETosis, stained for Ly6G and Ly6C, and analyzed for SYTOX green positivity on an Amnis CellStream flow cytometer (Cytek). Samples were run in technical duplicate. NETosis induction was calculated by normalizing the percent of SYTOX+ neutrophils (Ly6C^mid^Ly6G^+^ cells) to the average of percent SYTOX+ neutrophils in base media conditions (BMDNs incubated in 2% FBS/DMEM).

### Neutrophil-endothelial cell co-cultures

At 2 days prior to each timepoint of interest, irradiated and control ECs were collected and re-plated in 6-well plates. Cells were then incubated overnight and neutrophils (differentiated MPROs) were added in a 0.4µm Transwell insert (Corning) at a concentration of 1x10^6^ cells/mL in DMEM/2% FBS. Co-cultures were then incubated for 48 hours before collection for RNA isolation or western blot analysis.

### Western blot

Cell lysates were collected using RIPA buffer with 5mM EDTA, 1 mini cOmplete EASYpack tablet (Roche), and 1 mini PhoSTOP EASYpack tablet (Roche). Lysates were incubated on ice for 30 minutes under constant agitation before sonicating and centrifuging to pellet debris. Supernatant was then collected and stored at -80°C until use. To determine protein concentration, a bicinchoninic acid (BCA) assay (Pierce) was performed according to the manufacturer’s instructions. Lysates were denatured in 4x protein loading buffer (LICOR) with 10% β-mercaptoethanol and heated at 99°C for 10 minutes then cooled. Samples and a Precision Plus Protein Kaleidoscope Prestained Protein Ladder (Bio-Rad) were loaded into 4-20% gradient acrylamide gels with equivalent levels of total protein across all lanes. Gel electrophoresis was performed with a Bio-Rad PowerPac HC and mini-PROTEAN Tetra system and then protein transfer was performed overnight at 34V. Following transfer, membranes were dried, re-activated in methanol, and blocked for 1 hour in Intercept TBS buffer (LICOR). Membranes were then incubated with anti-phospho-p65 (Cell Signaling) and a loading control and incubated overnight at 4°C with constant agitation. Secondary antibodies were then incubated for 1 hour at room temperature under constant agitation before washing and reading on an Odyssey Fc Imager (LICOR). Analysis to determine protein expression relative to loading controls was performed in Fiji.

### Transwell endothelial monolayer model

C166 ECs were harvested and 75,000 cells were plated onto 3µm Transwell inserts (Corning). Inserts were incubated at room temperature for 20 minutes to enable even attachment prior to incubation at 37°C. Inserts were then cultured in complete media for 2 days to allow monolayer formation, which was confirmed via immunocytochemistry staining for ZO-1 and FITC-dextran permeability assays. Inserts were then irradiated and media was exchanged for DMEM/2% before incubating for indicated timeframes post-RT.

### Neutrophil transendothelial migration and FITC-dextran permeability assay

At 24 hours post-RT, endothelial monolayer inserts were washed with PBS and neutrophils (differentiated MPRO) were added to the inserts in serum-free IMDM/0.05% BSA. The receiver well below each insert contained 10ng/mL of CXCL2 in serum-free IMDM/0.05% BSA to facilitate neutrophil migration through the monolayer. Inserts were then incubated for 3 hours to allow neutrophil transendothelial migration. Neutrophils were then removed and monolayers were rinsed with PBS and transferred to a clean well containing PBS. To measure permeability, 40kDa FITC-dextran was added at a concentration of 1mg/mL to each insert and monolayers were incubated in the dark at room temperature for 20 minutes. Inserts were removed from the wells and the wells were analyzed for FITC intensity using a Varioskan LUX plate reader. Samples were run in technical duplicate and a standard curve of FITC-dextran was created at each experiment. For the standard curve, fluorescence intensity and concentration were plotted on a log-log axis and a line of best fit was established. For calculation of monolayer permeability, the fluorescence intensity across technical replicates was averaged and adjusted to a blank well containing only PBS. The concentration of FITC-dextran in each well was then determined using the standard curve and apparent permeability was calculated using the following equation, where the *V_basolateral_* is the volume in the receiver well (0.750mL); *C_sample_* is the calculated FITC-dextran concentration for each sample; *A* is the membrane surface area of the Transwell insert (0.3cm^2^); *C_0_* is the starting concentration of FITC-dextran (1000µg/mL); and *t* is the incubation time (1200 seconds):

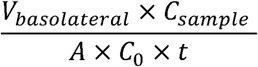

### siRNA treatment

Gene expression of *Rela* was knocked down using Silencer Select siRNA targeting *Rela* (Invitrogen) or Silencer Select Negative Control No. 1 (Invitrogen). Stock solutions of 10µM siRNA were prepared using RNase-free water. Two days prior to irradiation, C166 ECs were treated for 48h with complete media supplemented with 1nM siRNA and 1µL Lipofectamine RNAiMAX reagent (Invitrogen). Cells were then exposed to RT and processed accordingly for downstream assays. For Transwell monolayer siRNA experiments, cells were reverse transfected with the siRNAs at the time of plating and cultured for 48h to allow monolayer formation and transfection. Monolayers were then irradiated and processed for downstream analysis.

### Statistical analysis

All plots were made using GraphPad Prism. Imaging analysis was performed using Fiji (ImageJ). Statistical significance was determined via GraphPad Prism. Relevant statistical tests are indicated in figure legends.

## Results

### The recurrence-permissive niche is characterized by excessive vascular remodeling and requires neutrophils for tumor cell colonization

To study the association between low lymphocyte counts and recurrence, we used a model of radiation-induced recurrence, where systemic CD8^+^ T cell reduction induces tumor cell colonization of irradiated mammary tissue at 10 days post-RT^6^. In this model, female BALB/c mice were inoculated orthotopically with 4T1^luc^ TNBC cells, which served as a source of circulating tumor cells (CTCs), and the contralateral mammary fat pad (MFP) was irradiated to 20 Gy (**Figure 1a**). This dose is consistent with both intra- and post-operative RT in breast cancer patients and appropriate for studying normal tissue damage on an acute timescale^6,21^. In mice with CD8^+^ T cell reduction, we observed sustained neutrophil infiltration of irradiated tissue at 10 days post-RT (**Figure 1b-e**). This was not observed in unirradiated nor immunocompetent IgG control mice, establishing a link between prolonged neutrophil infiltration and tumor cell colonization. To validate this observation, we used an independent model of low lymphocyte counts by inoculating athymic Nu/Nu mice with 4T1^luc^ cells and irradiating the contralateral MFP as in our BALB/c model (**Figure 1f**). Neutrophils were also present in the MFPs of irradiated Nu/Nu mice at 10 days post-RT (**Figure 1g**), further demonstrating that neutrophils persist in the irradiated mammary of lymphocyte-deficient microenvironments.

**Figure 1.**
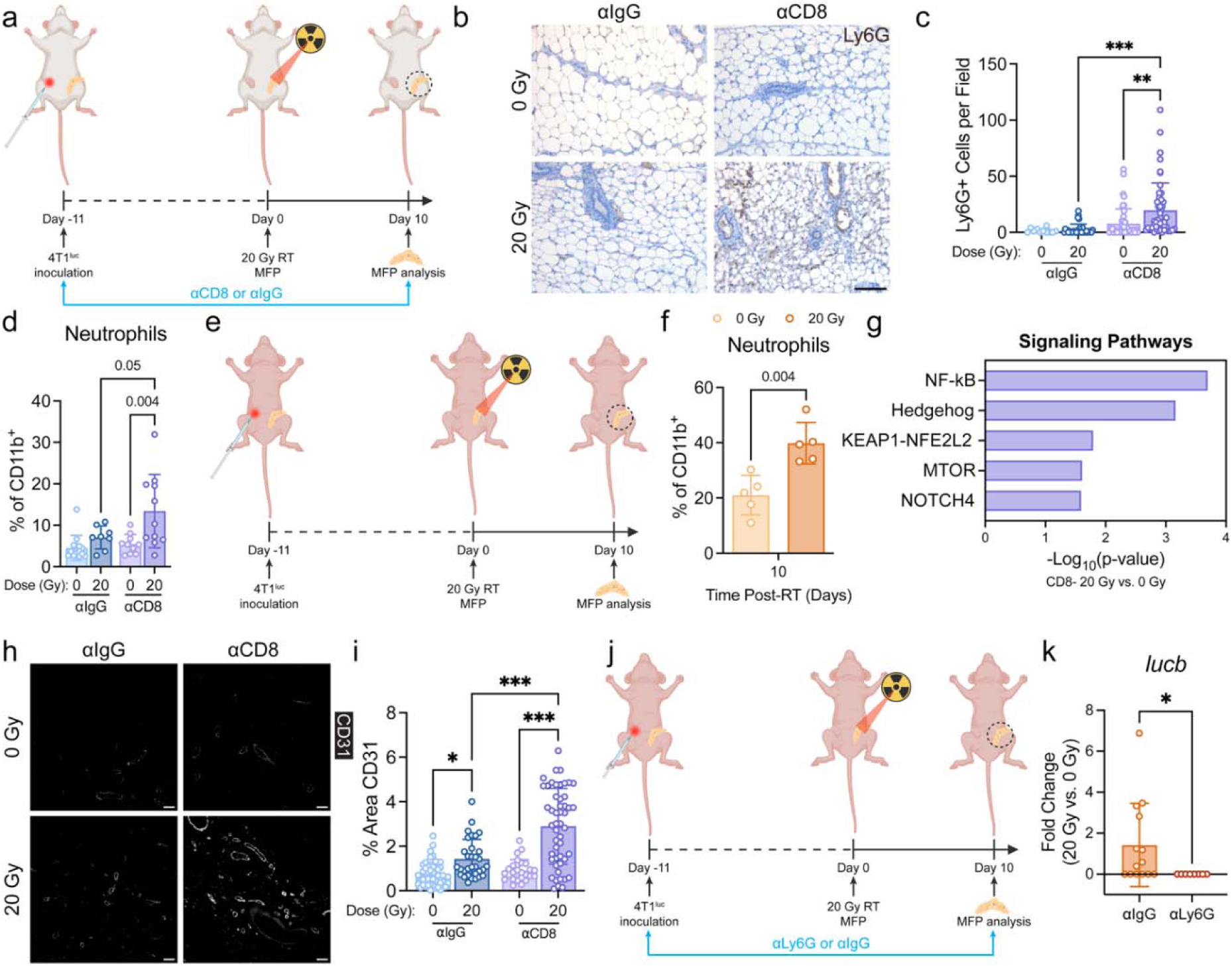
Neutrophils are required for tumor cell colonization of irradiated mammary tissue. **a.** Experimental timeline for CD8^+^ T cell redution experiments. Female BALB/c mice were inoculated with 4T1^luc^ cells and treated with 500µg of CD8 or IgG control antibodies at the time of inoculation and every 5 days therafter. When tumors were 100mm^3^, the contralateral MFP was irradiate to 20 Gy and MFPs were collected for downstream analysis at 10 days post-RT. Representative images **(b)** and quantification **(c)** of Ly6G^+^ cells in MFPs of BALB/c mice. n=5 mice per condition. **p<0.01 and ***p<0.001 by ANOVA with Sidak’s test. **d.** Quantification of CD45^+^CD11b^+^Ly6C^mid^Ly6G^+^SSC^hi^ neutrophils in MFPs at 10d post-RT analyzed via flow cytometry. n=8-12 mice per condition. p-values determined by ANOVA with Sidak’s test. **e.** Experimental timeline for Nu/Nu experiments with details as noted above except antibody treatments. **f.** Quantification of CD45^+^CD11b^+^Ly6C^mid^Ly6G^+^ neutrophils in MFPs of Nu/Nu mice at 10d post-RT analyzed via flow cytometry. n=5 mice per condition. p-value determined by t-test. **(g)** Significantly enriched pathways related to tissue remodeling in irradiated CD8 mice determined by Reactome analysis of MFP bulk RNA-seq data. n=5 mice per condition. **h.** Representative images **(b)** and quantification **(c)** of percent CD31^+^ area in MFPs of BALB/c mice. n=5 mice per condition. *p<0.05 and ***p<0.001 by ANOVA with Sidak’s test. **j.** Experimental timeline for neutrophil depletion model. Nu/Nu mice were inoculated and treated as in **(e)** with additional i.p. injections with Ly6G or IgG control antibodies at the time of inoculation and every other day thereafter. **k.** Fold change of *lucb* expression relative to *Actb* in 20 Gy vs. 0 Gy contralateral MFPs via RT-qPCR. n=8-12 mice per condition. p-value determined by Mann-Whitney test.

Recent pre-clinical studies of breast cancer and chronic sterile inflammatory conditions implicate neutrophil-mediated tissue remodeling in pre-metastatic niche formation and disease progression^16,22,23^. We performed RNA-sequencing of the MFPs from our BALB/c model to understand bulk microenvironmental tissue changes associated with the sustained neutrophil presence and identified predicted activation of several pathways associated with tissue remodeling, most notably NF-κB signaling which can be a critical regulator of inflammation and downstream angiogenesis (**Figure 1h**). Accordingly, we observed over a three-fold increase in the number of CD31^+^ vessels post-RT in mice with CD8^+^ T cell reduction (**Figure 1i-j**). While IgG control mice also exhibited a significant increase in vessel counts post-RT, the magnitude was significantly less than the CD8^+^ T cell reduced mice and can be attributed to some level of tissue remodeling and vascular repair observed in irradiated tissues.

Given these concurrent observations of sustained neutrophil presence, excessive vascular remodeling, and tumor cell colonization at 10 days post-RT in lymphocyte-deficient mice, we hypothesized that neutrophils could be a critical driver of the pro-tumor niche and subsequent colonization. Using anti-Ly6G to deplete neutrophils in our Nu/Nu model (**Figure 1k; Supplemental Figure 1a-b**), we quantified tumor cell colonization through *lucb* gene expression analysis at 10 days post-RT. Strikingly, we observed no detectable *lucb* expression in MFPs of neutrophil-depleted mice, suggesting that neutrophils are required for tumor cell colonization of the irradiated mammary tissue. Together, these findings demonstrate that RT is associated with sustained neutrophil infiltration and excessive vascular remodeling in lymphocyte-deficient microenvironments and establish neutrophils as a critical component of tumor cell colonization of mammary tissue.

### Pathologically activated neutrophils regulate vascular and immune responses in irradiated mammary tissue

Having established neutrophils as a functional regulator of tumor cell colonization of irradiated MFPs, we next sought to understand how they alter the microenvironment to drive this colonization. Histological analysis of the vasculature in neutrophil-depleted MFPs revealed that the excessive vascular remodeling that we associated with tumor cell colonization does not occur in the absence of neutrophils (**Figure 2a-b**). Importantly, the neutrophil-rich microenvironment was also characterized by a significant increase in VEGF expression localized within CD31^+^ vessels post-RT in contrast to neutrophil-depleted MFPs, which exhibited no change in VEGF post-RT (**Figure 2a, c**). In previous studies, we have also demonstrated that macrophages are required for tumor cell colonization of lymphocyte-reduced, irradiated mammary tissue^6,8^. Interestingly, neutrophil depletion abated the persistent macrophage infiltration that we observe up to 10 days post-RT (**Figure 2d-e**), suggesting a potential dependency of tumor cell colonization on neutrophil-macrophage interactions.

**Figure 2.**
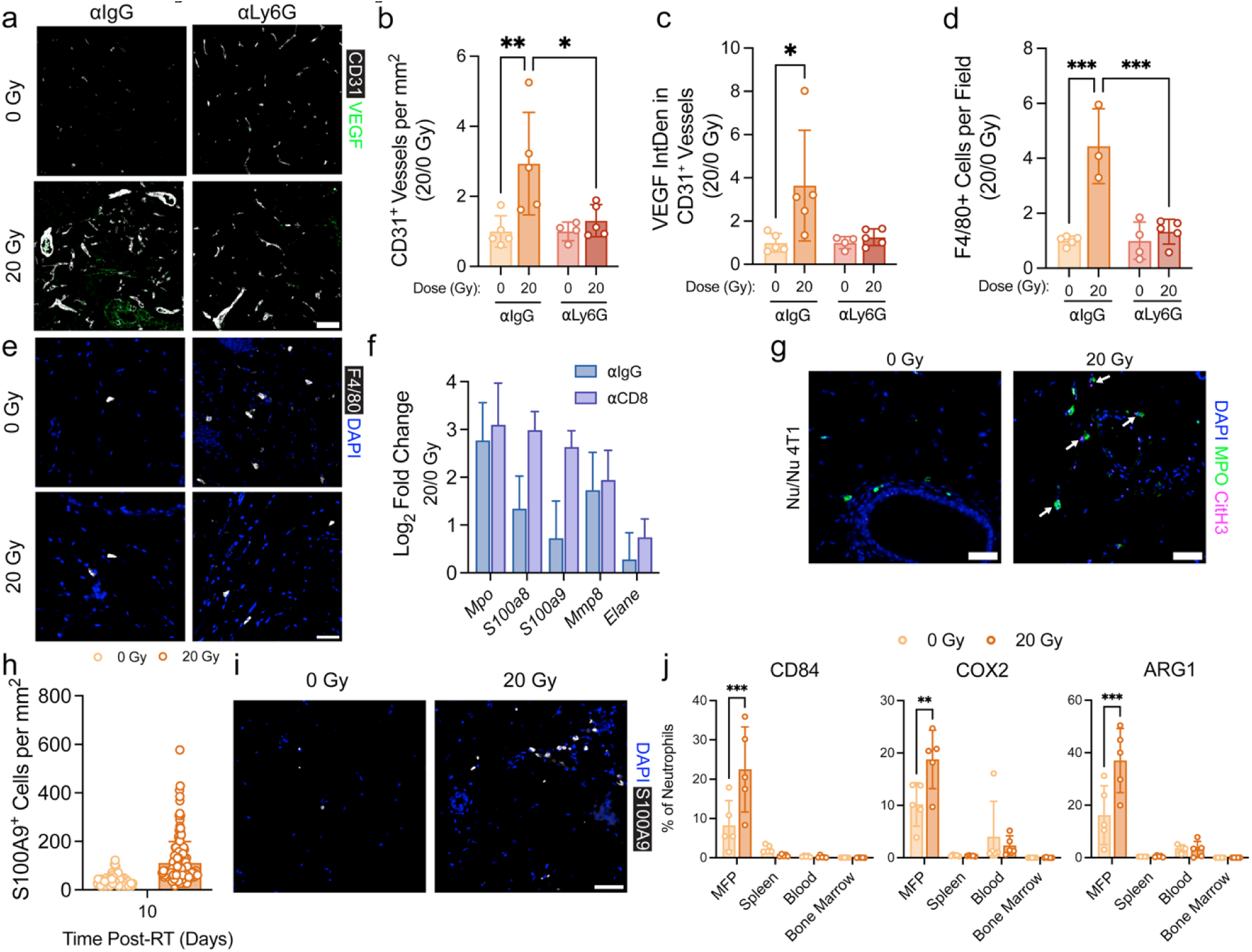
Pathologically activated neutrophils regulate the stromal and myeloid cell compartments of the pro-recurrent niche in irradiated mammary tissue. **a.** Representative images of CD31-VEGF costaining of MFPs in Nu/Nu neutrophil depletion model. Scale bars are 50µm. Coresponding quantification of CD31^+^ vessels per mm^2^ (b) and integrated density of VEGF localized with CD31^+^ vessels (c). n=4-5 mice per condition. *p<0.05 and **p<0.01 determined by two-way ANOVA with Sidak’s test. Quantification (d) and (e) representative images of F4/80^+^ cells per mm^2^ in MFPs from Nu/Nu neutrophil depletion model. n=3-5 mice per condition. ***p<0.001 determined by two-way ANOVA with Sidak’s test. f. Gene expression of select genes related to a pathologically-activated PMN-MDSC phenotype from bulk RNA-seq analysis of MFPs of IgG and CD8 mice in 20 vs. 0 Gy MFPs. n=5 mice per condition. g. Representative images of neutrophil extracellular traps (NETs) via localization of DAPI, MPO, and CitH3 in MFPs of Nu/Nu mice bearing contralateral 4T1^luc^ tumors. Arrows indicate localization of all three stains. Quantification (h) and representative images (i) of S100A9^+^ cells in MFPs from Nu/Nu-4T1^luc^ model. n=2-5 mice per condition j. Flow cytometry analysis of expression of key PMN-MDSC markers shown as percent of neutrophils. Neutrophils were defined as live CD45^+^Ly6C^mid^Ly6G^+^ cells in MFP, spleen, blood, and bone marrow from Nu/Nu 4T1^luc^ model. n=5 mice per condition. **p<0.01 and ***p<0.001 by two-way ANOVA with organ as the repeated measures factor followed by Sidak’s test. All data show mean±SD. All error bars are 50µm.

The ability of these neutrophils to regulate the mammary microenvironment and permit tumor cell colonization suggested that there may be a functional shift post-RT in lymphocyte-deficient mice. Growing evidence demonstrates that in the context of cancer, the functional and phenotypic heterogeneity of neutrophils can be ascribed to a spectrum ranging from classically activated neutrophils with acute inflammatory and antimicrobial functions to pathologically activated neutrophils, or polymorphonuclear myeloid-derived suppressor cells (PMN-MDSCs), with sustained immunosuppressive and tissue remodeling properties^11^. We therefore sought to characterize the neutrophils in the irradiated MFPs to identify whether they undergo pathological activation towards a PMN-MDSC phenotype. In CD8^+^ T cell-reduced BALB/c MFPs, we observed upregulation of several genes (*Mpo, S100a8, S100a9, Mmp8, Elane*) associated with PMN-MDSCs post-RT^24^. Immunocompetent controls also exhibited similar increases in *Mpo* and *Mmp8* compared to the CD8-reduced mice, suggesting that some neutrophils may be active in remodeling that tissue; however, the proportion of neutrophils in the mammary tissue remained close to baseline levels by this time (**Figure 1a**), likely reducing their impact compared to the immunocompromised mice. In our Nu/Nu model, we also observed expression of myeloperoxidase (MPO) and increased expression of S100A9 in irradiated MFPs (**Figure 2g-i**). We found that the MPO was often localized with citrullinated histone H3 (CitH3), which is a marker of NETosis and can be an immunosuppressive, pro-tumor function of pathologically activated PMN-MDSCs. Neutrophils in irradiated Nu/Nu MFPs also expressed increased levels of CD84, COX2, and ARG1 (**Figure 2j**), demonstrating a shift towards a PMN-MDSC phenotype and the immunosuppressive potential of these cells^25–27^. Importantly, this shift was only observed in the irradiated MFP and not in blood, spleen, nor bone marrow, highlighting that pathological activation of neutrophils occurs from the irradiated mammary microenvironment.

### The senescence-associated endothelial cell secretome pathologically activates neutrophils

The persistence of a pathologically activated neutrophil population at 10 days post-RT suggested that changes within the mammary adipose microenvironment may drive this activation and sustain neutrophils past their conventional, acute response window. Radiation is known to induce senescence and often a chronic senescence-associated secretory phenotype (SASP) characterized by stable cell cycle arrest and an inflammatory secretome capable of regulating the behavior of surrounding cells. Given our observation of neutrophil-regulated vascular remodeling in the pro-recurrent mammary adipose niche, we next investigated whether the vasculature associated with neutrophil infiltration could be undergoing senescence and contributing to the neutrophil phenotypic shift. We observed p21 expression in CD31^+^ vessels of irradiated MFPs from Nu/Nu mice in contrast to unirradiated MFPs where we saw little to no p21 expression (**Figure 3a**), suggesting that the ECs lining vessels in our model undergo cell cycle arrest at 10 days post-RT. To investigate effects of senescence *in vitro,* we used C166 murine ECs and characterized their RT-associated senescence response to establish a model for endothelial-specific senescence (**Figure 3b**). ECs underwent a significant reduction in EdU incorporation at 1 day post-RT that was sustained up to at least 3 days post-RT (**Figure 3c**). Control ECs also underwent a gradual reduction in EdU incorporation, likely due to the reduction in serum used to culture cells post-RT (see *Methods*). ECs also exhibited significantly higher p21 expression and a visible increase in senescence-associated beta-galactosidase (SA-β-gal) activity by 7 days post-RT (**Figure 3e-f**), signifying an active metabolic state despite the stable cell cycle arrest and lack of proliferation—a hallmark of senescent cells. Finally, we characterized the secretome of these ECs to understand whether their senescence could be associated with a secretory phenotype. As early as 2 days post-RT, we observed significant upregulation of key inflammatory SASP factors (IL-6, CXCL1, CCL2, VEGF) in the conditioned media of irradiated ECs (**Figure 3g**). Expression of these cytokines tended to increase over time, especially IL-6 which was upregulated over 10-fold compared to control ECs by 7 days, supporting evidence of an acquired SASP that becomes more persistent with time post-RT. We also observed significant upregulation of IL-6, TNF-α, and IL-1α in MFP homogenates post-RT (**Figure 3h**). Interestingly, this increase was only observed in mice with CD8^+^ T cell reduction, suggesting a potential SASP that is specific to the pro-recurrent niche.

**Figure 3.**
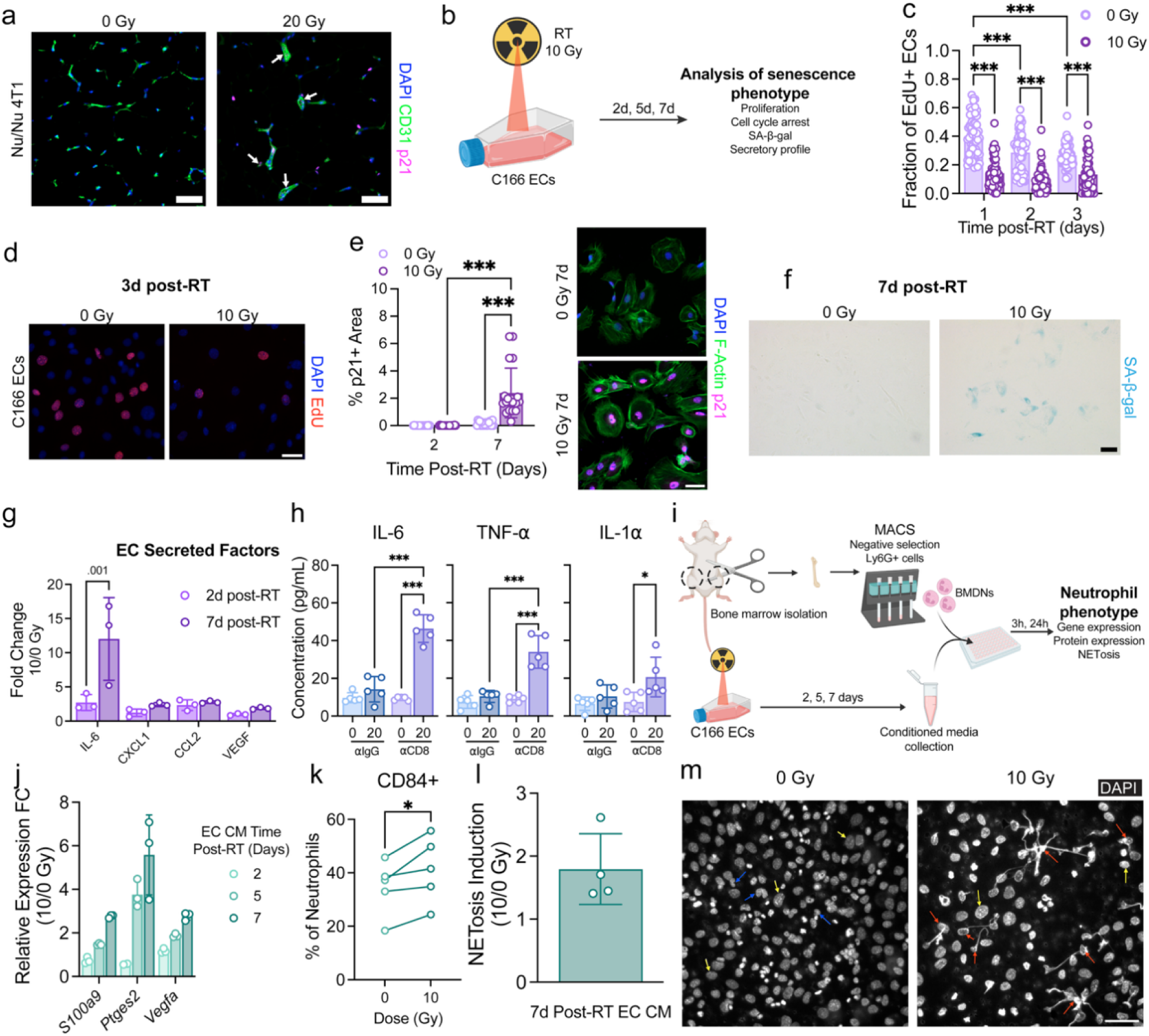
Senescent endothelial cells shift neutrophils toward a pathologically active state. **a.** Representative images of CD31/p21 costaining of MFPs from Nu/Nu model at 10 days post-RT. Arrows indicate examples of vessels (CD31) expressing p21. **b.** Experimental timeline for *in vitro* endothelial cell experiments. C166 ECs were irradiated to 10 Gy and cultured for 2, 5, or 7 days thereafter before analysis for hallmarks of senescence. Quantification **(c)** and representative images **(d)** of EdU incorporation in ECs post-RT. n=3 independent replicates per condition. **e.** Quantification of percent positive p21 staining in and representative images of ECs post-RT. n=3 independent replicates per condition. ***p<0.001 determined by ANOVA with Sidak’s. **f.** Representative images of SA-β-gal activity in ECs at 7 days post-RT. **g.** Cytokine profiling of EC conditioned media at 2 and 7 days post-RT. n=3 independent replicates per condition. Fold change calculated with respect to the average cytokine concentration of the 0 Gy sample at each corresponding timepoint. P-value determined by ANOVA with Sidak’s. **h.** Significantly upregulated SASP cytokines in MFP homogenate from BALB/c mice at 10 days post-RT. n=5 mice per condition. *p<0.05 and ***p<0.001 determined by ANOVA with Sidak’s. **i.** Experimental timeline of BMDN isolation and subsequent incubation in EC conditioned media. **j.** Gene expression of PMN-MDSC-related genes relative to Actb in BMDNs after incubation in EC CM. Error bars show SD. Quantification via flow cytometry of CD84 expression **(k)** and NETosis induction (SYTOX+ cells) **(l)** in Ly6G^+^Ly6C^mid^ BMDNs after incubation in EC conditioned media. n=5 and 2 independent experiments for NETosis and CD84 analysis, respectively. For CD84 analysis, p-values determined by paired t-test. **m.** Representative images of DAPI-stained irradiated (10 Gy) and control EC monolayers after neutrophil co-culture. Arrows show examples of EC (yellow) and neutrophil (blue) nuclei and activated neutrophils (red). All scale bars are 50 μm.

Despite their lack of cell division, senescent endothelial cells (ECs) have been demonstrated to contribute to concurrent vascular remodeling and regrowth through activation of neutrophils^22^. We hypothesized that radiation-induced senescence could initiate a positive feedback loop of unchecked inflammation and vascular remodeling wherein the EC SASP pathologically activates neutrophils, which then activate pro-remodeling molecular signaling in the ECs. As a first step in this feedback loop, we determined the effect of the EC SASP on neutrophils by culturing bone marrow-derived neutrophils in EC conditioned media and analyzing the neutrophils for pathological activation and a shift towards PMN-MDSCs (**Figure 3i**). BMDNs displayed upregulation of genes associated with the immunosuppressive, pro-tumor functions of PMN-MDSCs, including *S100a9*, *Ptges2*, and *Vegfa* (**Figure 3j**). We observed stepwise increases in expression of these genes as the BMDNs were cultured in conditioned media collected at increasingly longer timepoints, suggesting that the neutrophils become more pathologically activated as the EC SASP becomes more stable. We also observed increased expression of CD84 in neutrophils that were cultured in 7d irradiated conditioned media compared to control (**Figure 3k**). Finally, both indirect exposure from conditioned media and direct co-culture of neutrophils with ECs induced NETosis (**Figure 3m**). Taken together, these data support our hypothesis that the EC SASP can pathologically activate neutrophils in a manner consistent with our *in vivo* neutrophil phenotyping.

### Neutrophils amplify vascular dysfunction in irradiated ECs

To determine whether neutrophils activated by irradiated ECs could in turn activate molecular signaling and prolong dysfunction in the vasculature, we indirectly co-cultured neutrophils with ECs at acute and extended timepoints post-RT (**Figure 4a**). Given the role of NF-κB in inflammation and remodeling and its predicted upregulation in our *in vivo* model (**Figure 1h**), we analyzed the effect of neutrophils on induction of NF-κB signaling in irradiated ECs. ECs showed variable phosphorylated-p65 expression post-RT but displayed an increasing trend in expression when cultured with neutrophils as early as 6 hours post-RT (**Figure 4b**). At 7 days post-RT, we observed the largest increase in expression, with neutrophils inducing a 2-fold increase in phosphorylated-p65 expression in irradiated ECs, which was higher than the expression in ECs that received radiation alone. At the transcript level, we also saw that neutrophils increased expression of *Nfkb1* and *Nfkbia* at 7 days post-RT compared to ECs that were not co-cultured with neutrophils (**Figure 4c**), further suggesting ongoing activation of NF-κB signaling that is amplified by paracrine signaling with neutrophils. We also observed increases in downstream NF-κB genes, *Angpt2* and *Il1b* (**Figure 4d**), highlighting the potential of neutrophils to induce endothelial damage and exacerbate their inflammatory response.

**Figure 4.**
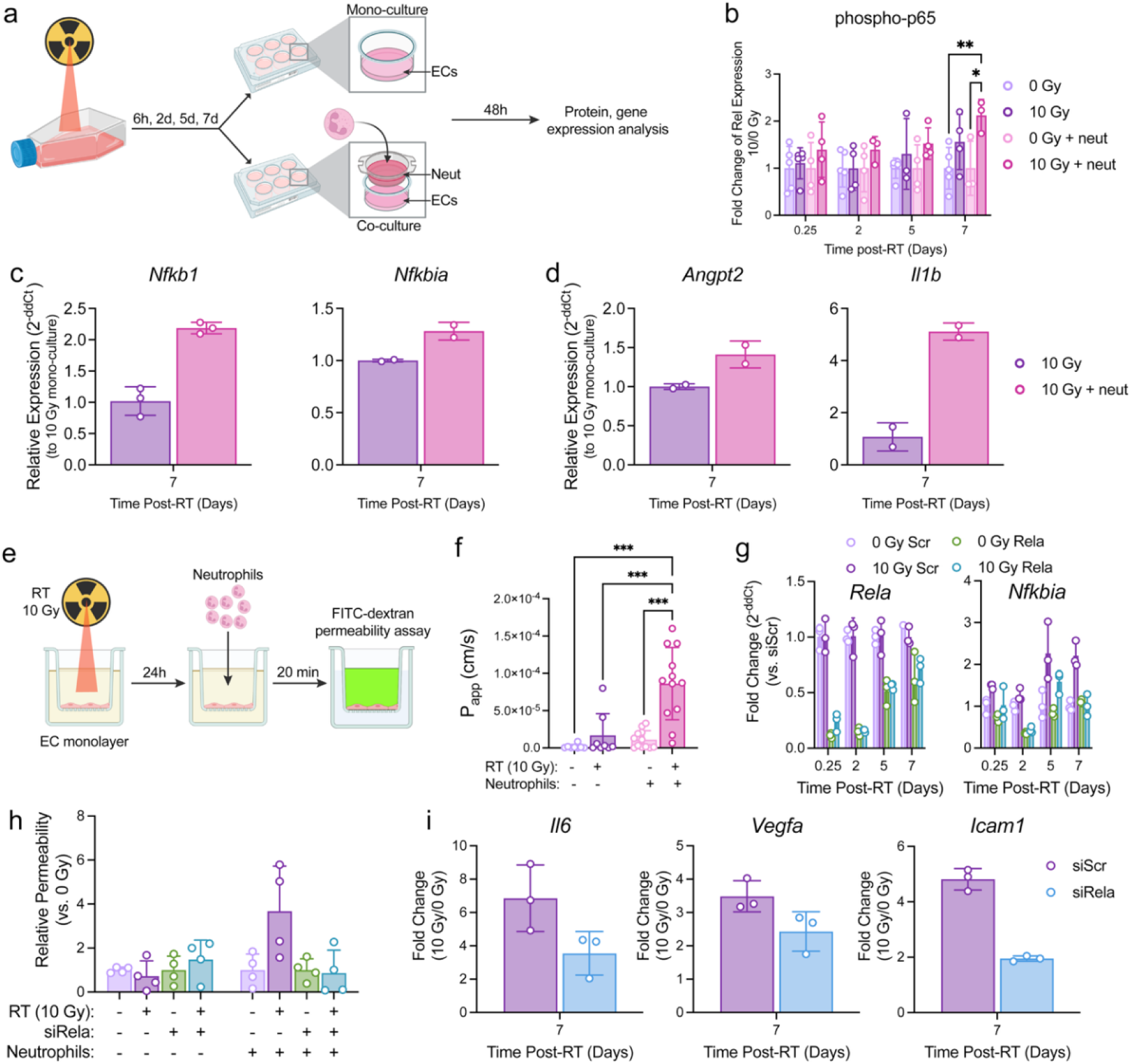
Endothelial dysfunction post-RT is exacerbated by neutrophils and NF-κB-dependent. **a.** Experimental timeline of neutrophil-EC co-culture experiments. ECs were irradiated to 10 Gy. At 24h prior to each timepoint of interest, co-cultures were initiated and cells were cultured for 48h thereafter before downstream gene and protein expression analysis. **b.** Western blot analysis of phosphorylated-p65 expression in ECs with and without neutrophil co-cultures. Protein expression was normalized to a loading control and fold change relative to each 0 Gy culture condition was calculated. n=3-5 independent experiments per condition. *p<0.05 and **p<0.01 by ANOVA with Sidak’s test. Gene expression analysis of genes related to NF-κB signaling **(c)** and vascular dysfunction/inflammation **(d)** in ECs with and without neutrophil co-culture. **e.** Experimental timeline for FITC-dextran permeability assays with EC Transwell monolayer model. Inserts were irradiated and cultured for 24h before performing neutrophil migration and subsequent FITC-dextran permeability assay. **f.** Quantification of apparent permeability of endothelial monolayers post-RT and post-neutrophil migration. n=5-12 independent experiments. ***p<0.001 by ANOVA with Sidak’s. **g.** Gene expression analysis to confirm knockdown of *Rela* and inhibition of downstream NF-κB signaling in irradiated si*Rela* and scramble control ECs up to 7 days post-RT. **h.** FITC-dextran permeability of si*Rela* and scramble control EC monlayers after RT and neutrophil migration. n=2 independent experiments. **i.** Gene expression analysis of si*Rela* and scramble control ECs post-RT.

Given these observations and the role of angiopoietin-2 in vascular permeability and inflammation, we next sought to determine the effect of neutrophils on endothelial monolayer permeability and determine whether neutrophils activated by irradiated ECs could exacerbate RT-induced vascular dysfunction. We used a Transwell model of an endothelial monolayer to directly co-culture neutrophils with irradiated ECs and measured changes in permeability using FITC-dextran (**Figure 4e**). Neutrophils significantly increased the permeability of irradiated endothelial monolayers post-RT, which exhibited a 5.2-fold increase in permeability compared to monolayers exposed to radiation alone (**Figure 4f**). To determine whether endothelial NF-κB signaling drives this neutrophil-induced increase in permeability, we used si*Rela* to prevent p65 signaling and analyzed the effects on irradiated ECs and subsequent neutrophil-induced permeability. We confirmed a sustained reduction in signaling up to 7 days post-RT with reduced expression of both *Rela* and *Nfkbia* in si*Rela* cells compared to scramble controls (**Figure 4g**). In our Transwell endothelial monolayer model, knockdown of *Rela* attenuated the neutrophil-induced increase in permeability observed in siScr ECs **(Figure 4h**). We also observed less induction of *Il6*, *Vegfa*, and *Icam1* in si*Rela* ECs at 7 days post-RT (**Figure 4i**), further highlighting the significance of *Rela* and canonical NF-κB signaling in the inflammatory, pro-remodeling endothelial cell response. Taken together, these findings demonstrate that in lymphocyte-deficient mammary tissue, RT establishes a persistent vascular inflammatory state that pathologically activates neutrophils, which further amplify NF-κB-dependent endothelial dysfunction and vascular remodeling to facilitate a pro-recurrent niche (**Figure 5**).

**Figure 5.**
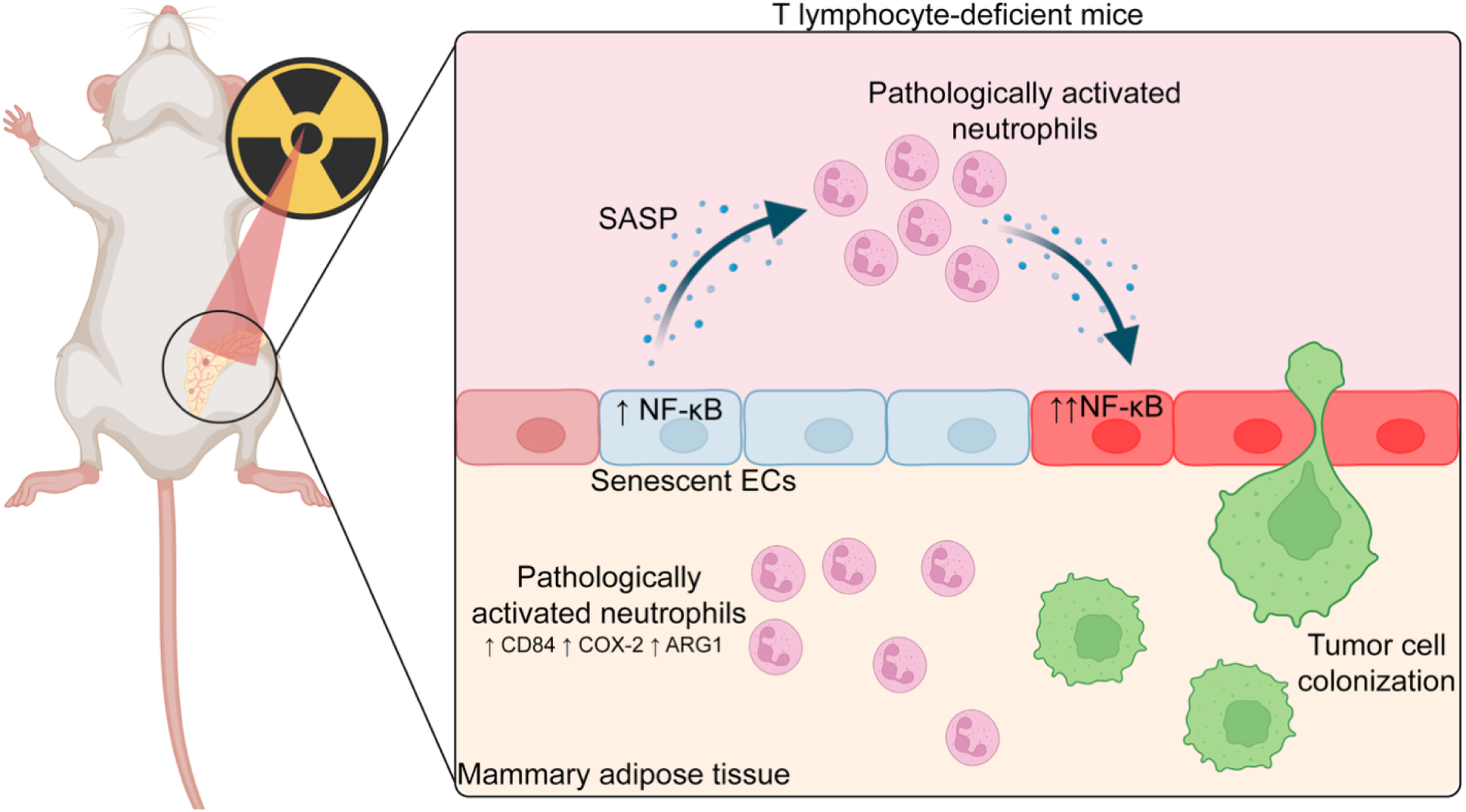
Hypothesized mechanism of a vascular-neutrophil feedback loop that drives tumor cell colonization. RT induces an NF-κB-dependent pro-inflammatory secretome from senescent ECs causes pathological neutrophil activation and subsequent amplification of NF-κB signaling, perpetuating a pathological neutrophil-EC feedback loop. This interaction may lead to tumor cell colonization in T lymphocyte-deficient mice, wherein RT induces a pathological neutrophil phenotype that coincides with neutrophil dependent vascular remodeling and tumor cell colonization.

## Discussion

RT is a critical component of TNBC treatment as it reduces the likelihood of recurrence^1^. Still, patients with chronic lymphopenia recur at significantly higher rates than patients with recovered lymphocyte counts^3,4^, representing a critical gap in our knowledge of how RT affects the breast microenvironment. We have previously shown in mouse models of lymphocyte deficiency that irradiation of the mammary adipose tissue initiates an immune cell-driven program where CTCs colonize the damaged normal tissue in the absence of T lymphocytes^6,8^. In the present study, we extended these observations to identify a potential link between high NLR—another predictor of outcome post-RT^7^—and neutrophil-induced changes to the irradiated microenvironment that make it more permissive to tumor cell colonization. We found that the pro-recurrent niche is characterized by neutrophil-dependent vascular remodeling and that neutrophils are required for tumor cell colonization of the irradiated mammary adipose tissue. It is well-established that systemic immune status can be linked to anti-tumor immunity^28–30^. Here, we provide further evidence that a patient’s systemic factors can also impact the normal tissue response to RT.

Our findings suggest that neutrophils regulate the broader mammary tissue response to RT rather than solely infiltrating the tissue as part of the conventional wound healing response to acute damage. Ly6G^+^ cell depletion attenuated the excessive vascular remodeling observed in lymphocyte-deficient tissue and prevented the persistent macrophage infiltration observed at 10 days post-RT. We previously showed that macrophages are also required for tumor cell colonization in lymphocyte-deficient mice^6^, suggesting the possibility that neutrophils may regulate an early stage of tissue remodeling that promotes subsequent macrophage recruitment or persistence. While additional investigation into the neutrophil-dependency of macrophage signaling in our model is required to draw conclusions, recent studies highlight the importance of neutrophils in macrophage-mediated tissue remodeling^31,32^, establishing a potential intermediate between neutrophils and our observed tissue remodeling. Taken together, the requirement of neutrophils for our observed excessive vascular remodeling, macrophage infiltration, and tumor cell colonization suggests that neutrophils contribute to a fundamentally altered tissue response to RT, providing a potential biological framework for why chronic systemic inflammation and a high NLR are associated with poor outcome.

Beyond the observed regulation by neutrophils of the pro-recurrent niche, we found that the neutrophils exhibited features associated with pathological activation. Neutrophils in the irradiated tissues of lymphocyte-deficient mice exhibited significantly increased levels of CD84, COX-2, S100A9, and ARG-1, as well as potential evidence of NETosis, in the irradiated MFPs. These features overlap with those identified in pathologically activated neutrophils and PMN-MDSCs that display increased immunosuppressive activity and pro-tumor function^14,26,27,33,34^. In addition to their immunosuppressive capacity, pathologically activated neutrophils have been implicated in tissue remodeling of pre-metastatic niches and in other chronic sterile inflammatory conditions^16,22,23^. While additional functional assays of immunosuppressive capacity and remodeling capability are required to further characterize the neutrophils in our model, our findings build upon the idea that neutrophils can persist in damaged tissues and drive chronic rewiring of microenvironments.

The persistence of an altered neutrophil phenotype beyond the initial acute phase of radiation injury and the lack of this phenotype in systemic immune compartments suggested microenvironment-specific activation of the neutrophils. Senescence is a known response of stromal cells to RT^35–37^ and as a gateway into the tissue, endothelial cells commonly undergo senescence as a stress response to protect the remaining tissue. We observed evidence of a SASP *in vivo* in the irradiated, lymphocyte-deficient tissue and further characterized a SASP in ECs *in vitro*. Both models were characterized by significant upregulation and secretion of pro-inflammatory cytokines, providing a potential link to why neutrophils persist in an activated state within the pro-recurrent niche. In fact, the secretome of senescent ECs recapitulated the pathological neutrophil activation *in vitro*, suggesting that ECs are an active regulator of radiation response in mammary tissue.

Our functional studies a reciprocal relationship between senescent ECs and pathologically activated neutrophils. We found that when indirectly co-cultured, neutrophils induce key indicators of canonical NF-κB signaling and increase the permeability of endothelial monolayers compared to RT alone. Upon knockdown of *Rela* in ECs, the neutrophil-induced increase in permeability post-RT was attenuated and ECs exhibited a reduction in vascular remodeling and inflammatory genes, including *Il6*, *Vegfa*, and *Icam1*. Taken together, these findings demonstrate NF-κB as a potential regulator of the reciprocal relationship between irradiated ECs and neutrophils, providing a framework for the persistent vascular dysfunction and pathological neutrophil activation. These findings are consistent with observations from our previous study where neutrophils exacerbated endothelial barrier permeability in a mammary vasculature-on-a-chip model^38^. In this model, we also observed subsequent increases in tumor cell adhesion to and spreading on the endothelium, suggesting that these interactions could be conducive to key steps of tumor cell tissue colonization. Thus, neutrophil-mediated barrier disruption may provide a mechanism of persistent RT-induced inflammation and vascular dysfunction that facilitate the early stages of tumor cell colonization.

While these studies establish a relationship between irradiated ECs and persistent neutrophil activation, the molecular signaling responsible for both endothelial and neutrophil signaling remain unclear. Our conditioned media studies demonstrate that endothelial soluble factors are sufficient for pathological activation of neutrophils, but the specific mediators of this response remain unknown. The SASP encompasses a diverse range of signaling molecules^35,39^, and it is likely that there are multiple factors contributing to our observed response. Still, investigation into key drivers of neutrophil phenotype shift would provide insight into potential targets to disrupt the EC-neutrophil feedback loop. Similarly, the neutrophil-derived factors responsible for activating phosphorylation of p65 in irradiated ECs and the functions responsible for increasing permeability of the endothelial monolayer have not yet been identified. We observed web-like structures on irradiated ECs in our Transwell model and increased NETosis induction of BMDNs exposed to the irradiated EC secretome. NETs can be decorated with various granule proteins and proteases that can induce EC death^22,40,41^, providing a potential mechanism of neutrophil-mediated endothelial damage consistent with previous findings; however, additional studies have demonstrated that neutrophil oxidative burst and reactive oxygen species also increase endothelial permeability^23^, thereby warranting further investigation. In addition to canonical NF-κB signaling, non-canonical signaling has been demonstrated to mediate tissue damage^42^. Investigation into the interplay of these signaling pathways is critical as non-canonical signaling provides a promising therapeutic target for chronic tissue remodeling that may exhibit less off-target effects that canonical signaling^43–45^. Finally, the functional requirement of endothelial NF-κB must be tested *in vivo* to determine the relevance for tumor cell colonization of the irradiated mammary adipose tissue.

## Conclusions

Identifying mechanisms of recurrence requires models with clinically relevant systemic conditions to understand why certain risk factors lead to recurrence. We identified significant differences in the irradiated mammary adipose microenvironment between mice with and without lymphocyte-deficiencies, further underscoring the impact of systemic immune status on TNBC outcomes. By establishing a hypothesized feedback loop between RT-damaged ECs and pathologically activated neutrophils, we developed a framework for future investigation into two potential points of intervention: RT-induced endothelial senescence and pathologically activated neutrophils. Given the associations of lymphopenia, chronic inflammation, and TNBC relapse post-RT, resolving these interactions on a molecular level could lead to development of recurrence interventions and significantly improve outcomes for high-risk TNBC patients.

## Acknowledgements

We thank Dr. Michael Freeman for irradiator use and Dr. Alissa Weaver for flow cytometer use. The Vanderbilt Flow Cytometry Shared Resource administers the FlowJo Portal Group License used to analyze the flow cytometry data and is supported by the Vanderbilt Ingram Cancer Center (P30CA068485) and the Vanderbilt Digestive Disease Research Center (DK058404). This research was financially supported by the Vanderbilt Institute for Clinical and Translational Research Grant Number VR74036, National Institutes of Health Grant Numbers R00CA201304 (M.R.), F31CA298725 (S.E.M), and T32DK101003 (K.C.), and a Vanderbilt Scaling Success Award. Schematics were created using BioRender.

## Supplemental Figures

**Figure S1.**
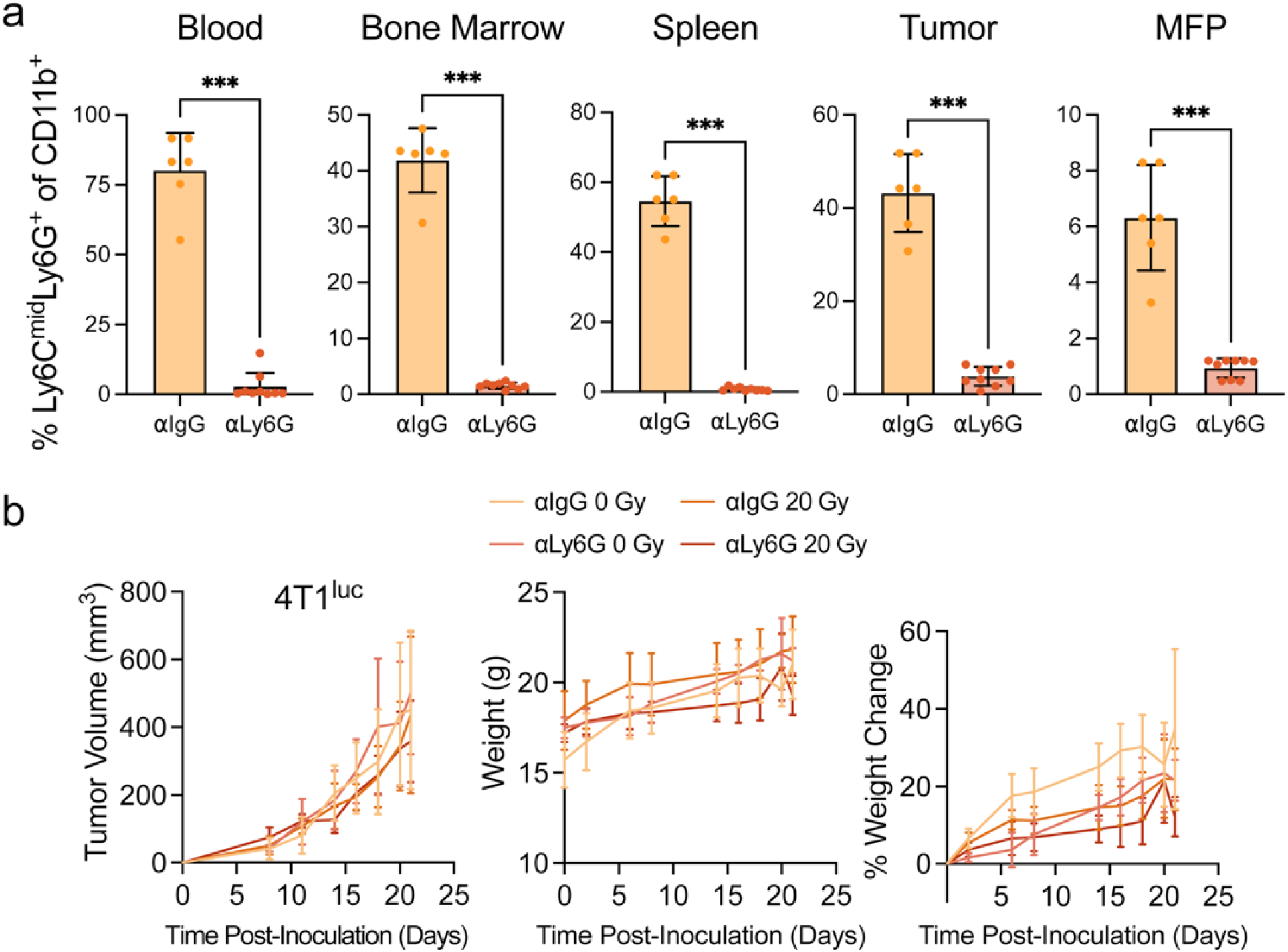
Confirmation of neutrophil depletion. **a.** Flow cytometry analysis of CD45^+^CD11b^+^Ly6C^mid^Ly6G^+^ cells in Nu/Nu-4T1^luc^ neutorphil depletion model without RT. n=6-9 mice per condition. ***p<0.001 by Mann-Whitney test. **b.** Mouse metrics over time in Nu/Nu-4T1^luc^ neutrophil depletion model with RT. n=5 mice per condition. Data show mean±SD.

